# Forced isoform switching of Neat1_1 to Neat1_2 leads to the loss of Neat1_1 and the hyperformation of paraspeckles but does not affect the development and growth of mice

**DOI:** 10.1101/698068

**Authors:** Momo Isobe, Hikaru Toya, Mari Mito, Tomoki Chiba, Hiroshi Asahara, Tetsuro Hirose, Shinichi Nakagawa

## Abstract

Neat1 is a long noncoding RNA (lncRNA) that serves as an architectural component of the nuclear bodies known as paraspeckles. Two isoforms of Neat1, the short isoform Neat1_1 and the long isoform Neat1_2, are generated from the same gene locus by alternative 3’ processing. Neat1_1 is the most abundant and the best conserved isoform expressed in various cell types, whereas Neat1_2 is expressed in a small population of particular cell types, including the tip cells of the intestinal epithelium. To investigate the physiological significance of isoform switching, we created mutant mice that solely expressed Neat1_2 by deleting the upstream polyadenylation (poly-A) signal (PAS) required for the production of Neat1_1. We observed the loss of Neat1_1 and strong upregulation of Neat1_2 in various tissues and cells and the subsequent hyperformation of paraspeckles, especially in cells that normally express Neat1_2. However, the mutant mice were born at the expected Mendelian ratios and did not exhibit obvious external and histological abnormalities. These observations suggested that the hyperformation of paraspeckles does not interfere with the development and growth of these animals under normal laboratory conditions.

## Introduction

Many long noncoding RNAs (lncRNA) are transcribed from the genome, and these transcripts regulate various molecular and cellular processes, including the epigenetic control of gene expression, mRNA processing, and the formation of nonmembranous cellular bodies (Long et al. 2017). Neat1 is one of the best-studied functionally validated lncRNAs and serves as an architectural component of the nuclear bodies known as paraspeckles (Chen and Carmichael 2009; Clemson et al. 2009; Sasaki et al. 2009; Sunwoo et al. 2009). Paraspeckles are phase-separated nonmembranous nuclear bodies that are unique to mammalian species (Fox et al. 2018; Nakagawa et al. 2018); more than 40 proteins assembled at a specific region of Neat1 (Fox et al. 2002; Dettwiler et al. 2004; Fox et al. 2005; Naganuma et al. 2012; Fong et al. 2013; Hu et al. 2015; Kawaguchi et al. 2015) form a characteristic core-shell structure (Souquere et al. 2010; West et al. 2016). Paraspeckles have been proposed to function via the sequestration of paraspeckle-associated RNA binding proteins within these nuclear bodies (Hirose et al. 2014; Imamura et al. 2014). Two isoforms of Neat1, Neat1_1 (3.2 kb in mouse and 3.7 kb in human) and Neat1_2 (20 kb in mouse and 23 kb in human), are generated from the same locus by alternative 3’ processing (Naganuma et al. 2012). Neat1_1 transcripts are processed by the canonical 3’ end processing complex CPSF6– NUDT21 (CFIm) and are cleaved at the polyadenylation (poly-A) signal (PAS) located upstream. The 3’ processing of Neat1_1 is inhibited by Hnrnpk, which binds to sequences between the CFIm binding site and the PAS, resulting in the generation of readthrough products that give rise to the long isoform Neat1_2 (Naganuma et al. 2012; Yamazaki et al. 2018). Alternative polyadenylation is also regulated by Tardbp, which binds to the GU-rich motifs upstream of the PAS (Modic et al. 2019). The 3’ end of Neat1_2 is cleaved by RNase P, and the non-polyadenylated transcripts are stabilized by triple-helix RNA structures found in the terminal region of Neat1_2 (Wilusz et al. 2012; Yamazaki et al. 2018). The specific knockdown of Neat1_2 leads to the disassembly of paraspeckles even in the presence of intact Neat1_1, suggesting that Neat1_2, but not Neat1_1, is required for the formation of paraspeckles (Sasaki et al. 2009). While most cultured cell lines express both Neat1_1 and Neat1_2 and form paraspeckles, strong expression of Neat1_2 is restricted to a subpopulation of particular cell types in mouse tissues, and Neat1_1 expression is widespread (Nakagawa et al. 2011). Accordingly, prominent paraspeckle formations are found in only the small population of cells expressing high levels of Neat1_2, including the tip cells of the intestinal epithelium, the surface mucous cells of the stomach, and the corpus luteal cells in the ovary (Nakagawa et al. 2011). Cells that express only Neat1_1 do not form paraspeckles, and Neat1_1 is diffusely localized to the nucleoplasm of these cells (Nakagawa et al. 2011; Li et al. 2017). Consistent with the strong expression of Neat1_2 in corpus luteal cells, the fertility of female Neat1 knockout (KO) mice is severely decreased due to the impaired formation of the corpus luteum and the subsequent decrease in serum progesterone (Nakagawa et al. 2014). Notably, only half of the Neat1 KO female mice exhibit a subfertile phenotype, and the corpus luteal cells exhibit normal differentiation in the fertile mutant mice, without the formation of paraspeckles. Neat1 KO mice also exhibit defects in the lactation (Standaert et al. 2014) and the phenotypic switching of vascular smooth muscle cells in response to vascular injury (Ahmed et al. 2018), suggesting that Neat1 is required for various cellular processes in physiological conditions. In addition, Neat1 can enhance or suppress cancer progression in a condition-specific manner (Adriaens et al. 2016; Mello et al. 2017). Thus, paraspeckles are proposed to become essential only when mice are placed in certain environments, of which the precise molecular conditions have not been identified (Nakagawa et al. 2011; Nakagawa et al. 2014).

In this study, we forced the isoform switching of Neat1 by deleting the PAS required for the production of Neat1_1, which has been shown to increase the expression of Neat1_2 at the expense of Neat1_1 expression (Li et al. 2017; Wang et al. 2018; Yamazaki et al. 2018; Modic et al. 2019). Despite the lack of Neat1_1 expression and hyperformation of paraspeckles in various tissues and cells, no overt phenotype was observed in the mutant mice lacking the Neat1 PAS 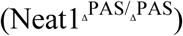. Our observation was consistent with the previous hypothesis that paraspeckles are dispensable under normal laboratory conditions and function only when cells are placed in particular conditions.

## Results

### Generation of mice that lack the PAS essential for the production of Neat1_1

To generate an animal model that displayed forced isoform switching from Neat1_1 to Neat1_2, we created mutant mice that lacked the PAS of Neat1 using CRISPR/Cas9-mediated genome editing. To test the cleavage efficiency of multiple gRNAs, we initially transfected 3T3 cells with plasmids that express Cas9 nuclease and each gRNA targeting the region surrounding the Neat1 PAS (**Figure 1A**). Genomic DNA was isolated from the cells, and DNA fragments containing the cleavage site were amplified by PCR and subjected to T7 endonuclease assays. Among the 5 gRNAs tested, one gRNA that cleaved the sequences inside the PAS (AATAAA) yielded the expected cleavage product at the highest efficiency (**Figure 1B**). We then injected the in vitro-transcribed gRNA and Cas9 mRNA into fertilized eggs and obtained 6 lines with various mutations (**Figure 1C**). After a single backcross to C57BL6/N wild-type (WT) mice, we examined the expression of Neat1_2 and Neat1_1 in the salivary glands of heterozygous mice. Because a mutant line that lacked 39 nucleotides, including the PAS, exhibited the highest expression of Neat1_2 relative to the expression of Neat1_1, we focused on this mutant allele 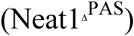 in the following analyses (**Figure 1D**). To confirm the expression of Neat1 isoforms in the 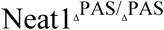 mice, we crossed heterozygous mice and prepared mouse embryonic fibroblast cells (MEFs) from WT and 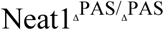 embryos. We then purified RNA from the MEFs and performed Northern blot analyses using probes that detected both Neat1_1 and Neat1_2 and probes that specifically detected Neat1_2 (**Figure 1F**). The expression of Neat1_1 was completely lost in the 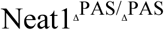 MEFs, whereas Neat1_2 was moderately upregulated in these cells (**Figure 1F**). The increase in Neat1_2 was also confirmed by in situ hybridization (**Figure 1G**) and resulted in enhanced paraspeckle formation, as shown by the accumulation of the paraspeckle marker protein Nono (**Figure 1G, H**). The 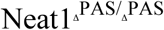 mice were born at the expected Mendelian ratio 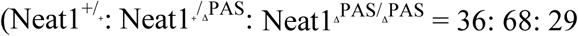 for males and 24: 54: 21 for females, p = 1.00 and 0.96, respectively, with a chi-squared test), exhibited a normal body weight 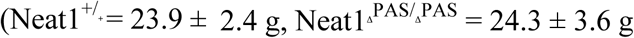 for males at 12–16 weeks, p = 0.89 with Student’s t-test), and had no overt morphological or behavioral defects.

**Figure 1.**
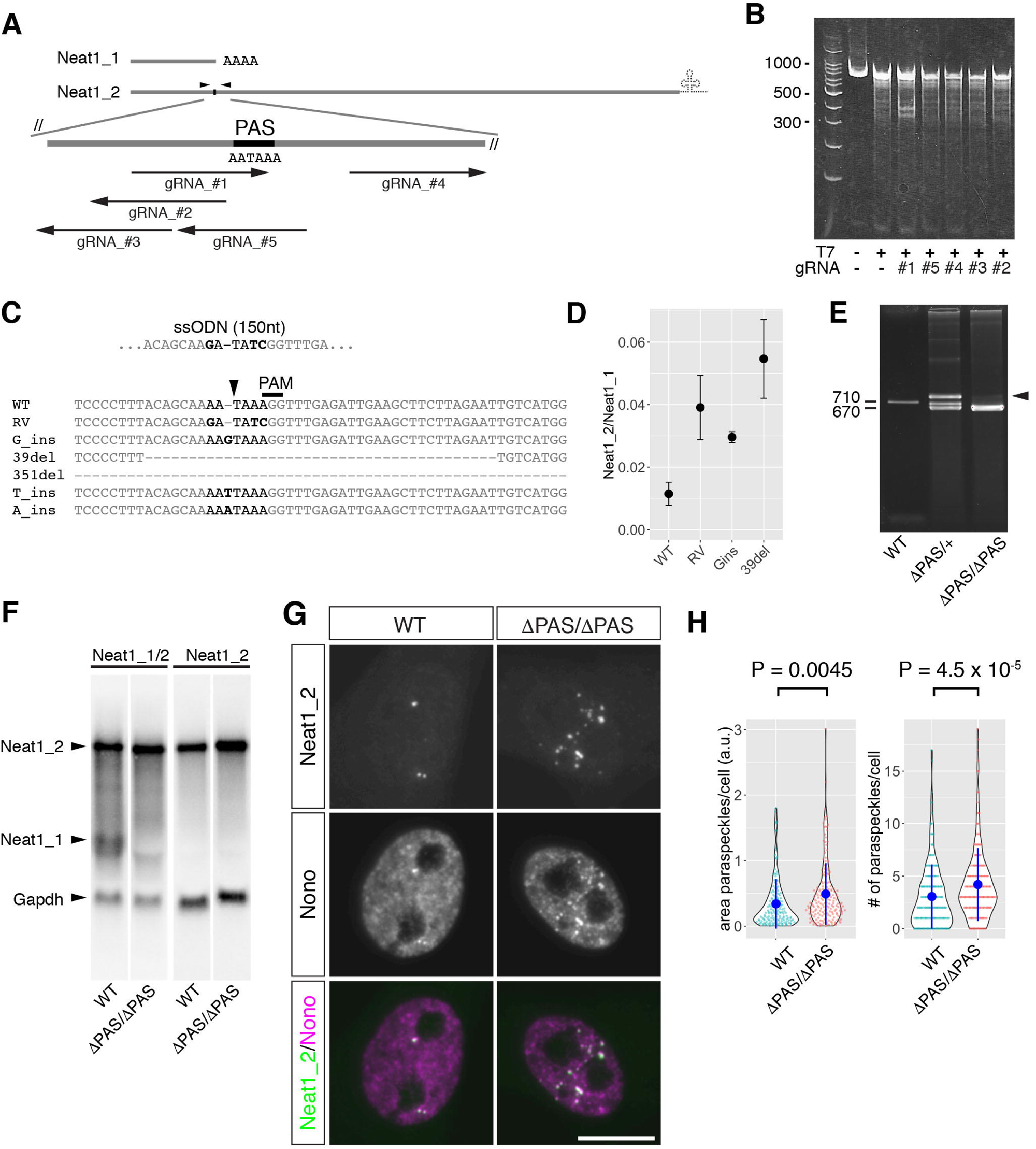
Generation of 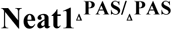 mice. (A) Schematics of gRNAs targeting the region surrounding the polyadenylation (poly-A) signal (PAS) AATAAA. Arrowheads indicate the positions of the primers used for the T7 endonuclease assay. (B) A T7 endonuclease assay evaluated the efficiency of gRNA-Cas9 endonuclease activity. Note that gRNA #1 exhibited the highest activity. (C) Sequence analysis of genome-edited mice. Six different mutant alleles were observed. The arrowhead indicates the position of the predicted cleavage site. Bold characters indicate the mutated nucleotides. (D) Comparison of the ratio of Neat1_1 and Neat1_2 expression with mutant alleles. Total RNA was obtained from the salivary glands of mutant mice (n = 2 for each line), and Neat1 expression was analyzed by RT-qPCR and calculated from technical duplicates. Note that 39del (ΔPAS) exhibited the highest Neat1_2/Neat1_1 ratio. p values were calculated by Student’s t-test. Blue dots and bars represent the mean value and the standard deviation for the biological duplicates, respectively. (E) An agarose electrophoresis gel image of PCR products that were amplified with genotyping primers. Note that a heteroduplex band that exhibited low mobility was obtained from a heterozygous mouse (arrowhead). (F) Northern blot analyses of total RNAs obtained from mouse embryonic fibroblast (MEF) cells derived from wild-type (WT) and 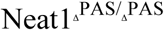 (ΔPAS/ΔPAS) mice. Neat1_1 was completely absent, whereas Neat1_2 was upregulated in 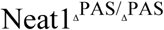 mice. (G) Simultaneous detection of Neat1_2 and Nono in MEF cells prepared from WT and ΔPAS embryos. Scale bar, 10 µm. Note that the number of paraspeckles increased in KO MEFs. (H) Quantification of the area and the number of paraspeckles shown in F, as visualized by violin/beeswarm plots. Two independent MEFs for each genotype were prepared from different embryos, and 100 cells for each batch of MEFs were randomly selected for the measurements. The values were mixed for each genotype and were used for the statistical analyses. p values were calculated by Wilcoxon’s rank sum tests. Blue dots and bars represent the mean value and the standard deviation for the biological replicates, respectively.

### Forced isoform switching by PAS deletion leads to the differential upregulation of Neat1_2 in each tissue in 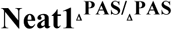 mice

To confirm the isoform switching of Neat1_1 and Neat1_2 in various tissues from WT and ΔPAS mice, we performed Northern blot analyses using RNA prepared from each tissue (**Figure 2, Supplemental Figure S1**). Three 16-week-old male mice were used for this study. In all cases, Neat1_1 expression was completely lost in 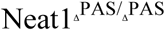 tissues, as well as MEFs (**Figure 2B, D**). However, the upregulation of Neat1_2 was variable in different tissues (**Figure 2C, D**). For example, brown adipose tissue and intestinal tissue exhibited greater fold changes in Neat1_2 (4.12- and 3.9-fold, respectively; **Figure 2D**), whereas the testis and salivary gland exhibited much smaller fold changes (1.88 and 1.82, respectively) than the appropriate WT tissues. This result was unexpected because the salivary gland expressed high levels of Neat1 and low levels of Neat1_2, and a forced isoform switch was anticipated to result in strong production of Neat1_2. We also found that the expression of Neat1_1 was almost absent in the liver (**Figure 2B**), which was not evident in previous in situ hybridization studies due to the lack of probes that specifically detected the short isoform (Nakagawa et al. 2011). The upregulation of Neat1_2 was detected with probes that specifically targeted the long isoform (**Figure 2C**). The upregulation of Neat1_2 appeared much weaker with probes that simultaneously detected Neat1_1 and Neat1_2 (**Figure 2B**) than with the Neat1_2 specific probes, probably because the nonspecific probe weakly cross-reacted with other transcripts that showed a similar size to Neat1_2 and were detected in the samples derived from Neat1 KO mice (**Supplemental Figure S2**). RT-qPCR analyses using primers specific for the Neat1_2 region also confirmed the variable upregulation of Neat1_2 in different tissues from 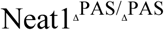 mice (**Supplemental Figure S3**).

**Figure 2.**
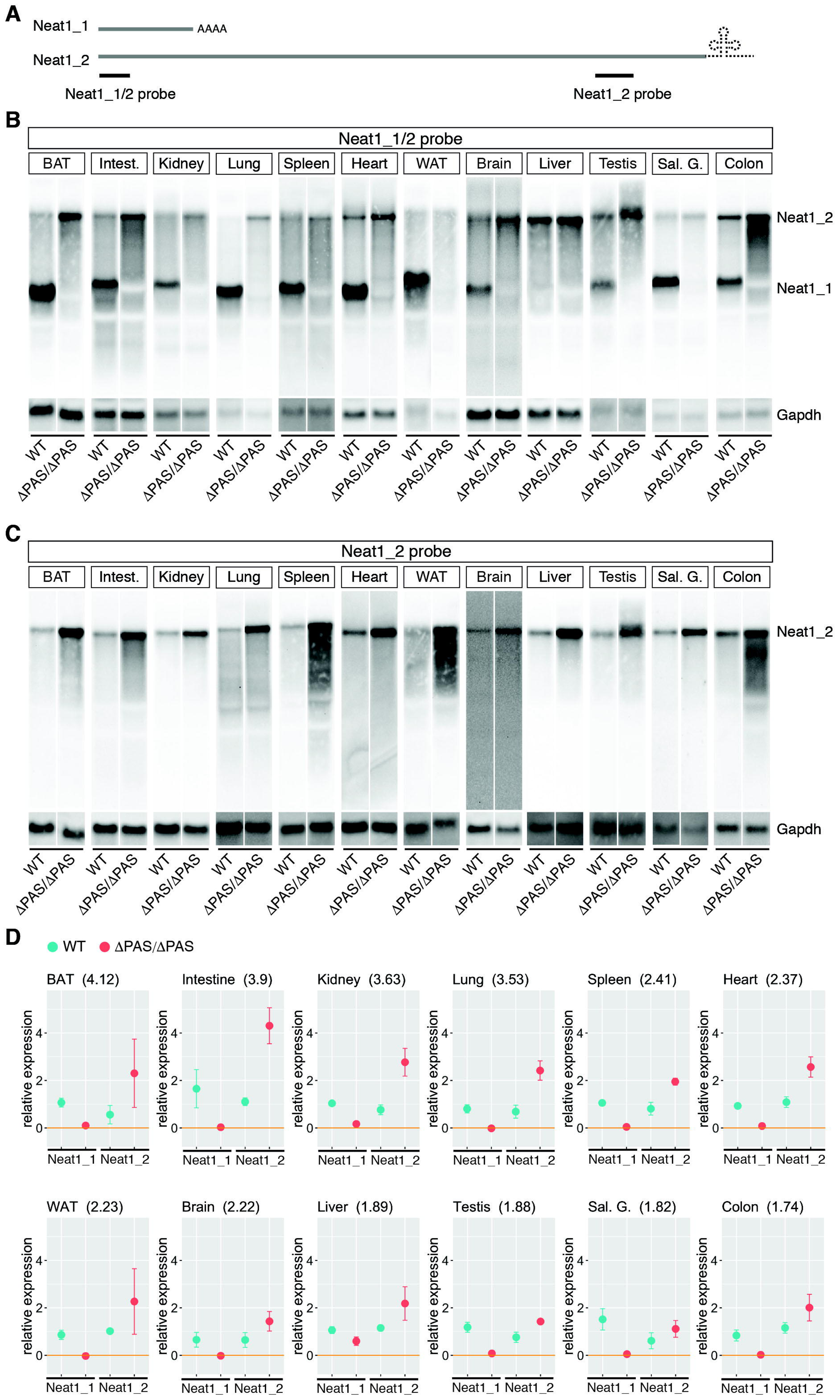
Northern blot analyses of Neat1 expression in various tissues of 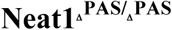 mice. (A) Schematics of the probes used for the detection of Neat1_1 and Neat1_2. The Neat1_1/2 probe detected both isoforms, whereas the Neat1_2 probe targeted a region specific to the long isoform. (B, C) Expression of the Neat1 isoforms, as revealed by probes that detected Neat1/2 (B) and Neat1_2 (C). Note the complete loss of Neat1_1 signals in all tissues and the variable upregulation of Neat1_2 in each tissue from 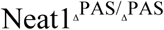 (ΔPAS/ΔPAS) mice. The Neat1_1/2 probe failed to detect the upregulation of Neat1_2 in certain tissues for unknown technical reasons. The liver expresses only Neat1_2, whereas the salivary gland expresses mostly Neat1_1. (D) Quantification of the results shown in B and C. The expression of Neat1_2 was quantified using the data shown in **Supplemental Figure S1**. The dots and bars represent the mean value and the standard deviation for the biological triplicates (duplicates for WT colon and white adipose tissue), respectively.

To study the isoform switching of Neat1 in the 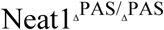 mice and to study paraspeckle formations at the cellular level, we examined the expression of Neat1_1/2 and Neat1_2 by in situ hybridization in three representative tissues, the intestine, liver, and salivary gland, in 8-week-old adult male mice (**Figures 3, 4**). In the intestine, strong expression of Neat1_2 was restricted to the tip cells of the intestinal epithelium (**Figure 3A**), as previously described (Nakagawa et al. 2011). The areas that expressed Neat1_2 were expanded and included the middle region of the intestinal epithelium in the 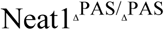 mice (**Figure 3A**). The ectopic formation of paraspeckles in the middle region of the intestinal epithelium was confirmed by the colocalization of the paraspeckle marker proteins Nono and Neat1_2 (**Figure 4A**). Paraspeckle formation also increased in the tip cells in the intestinal epithelium, which normally form paraspeckles (**Figure 4A**). We also observed enhanced expression of Neat1_2 and the subsequent hyperformation of paraspeckles in the liver (**Figure 3B, Figure 4A**), despite the lack of Neat1_1 expression observed in a Northern blot (see discussion). In the salivary glands of WT mice, almost no signals were observed with probes that detected Neat1_2, and intense signals were obtained with probes that detected Neat1_1/2 in granular duct cells, suggesting that these cells almost exclusively expressed Neat1_1 (insets in **Figure 3C**). The signals obtained with the Neat1_1/2 probe were substantially reduced in the 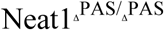 mice (**Figure 3C**), which was consistent with the absence of Neat1_1 transcript signals on a Northern blot (**Figure 2B**). Statistical analyses also confirmed the increases in the numbers and the areas of paraspeckles per cell in these tissues (**Figure 4B, C**), except for the area of paraspeckles in the middle regions of intestinal epithelium, where a small population of cells expressed Neat1_2 and formed prominent paraspeckles (note that the areas of paraspeckles were measured only for paraspeckle-containing cells and not for the cells that do not form paraspeckles). We examined the histology of the three tissues using hematoxylin and eosin (HE) staining, and no gross abnormalities were observed (**Figure 3A-C**). For these immunohistochemical and histochemical analyses, we examined 6 individual mice for each genotype, which exhibited consistent results.

**Figure 3.**
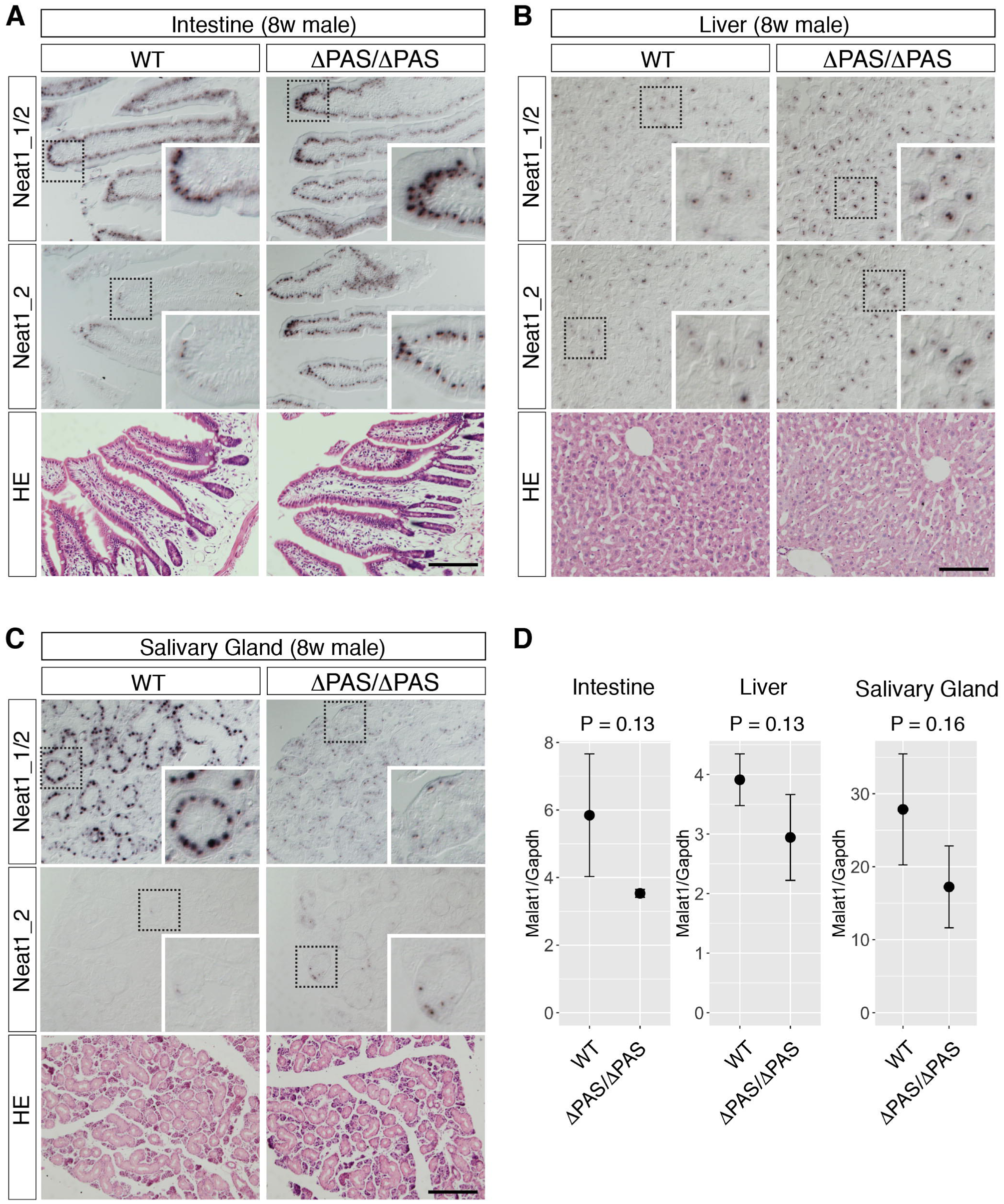
In situ hybridization analyses of Neat1 expression in 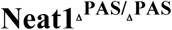 mice. (A-C) Expression of the Neat1 isoform revealed by probes that detected Neat1_1/2 and Neat1_2 in the intestine (A), liver (B), and salivary gland (C). Neat1_2 expression is normally restricted to the tip cells of the intestinal epithelium and expanded to the middle area of the epithelium in 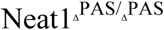 (ΔPAS/ΔPAS) mice. Signals detected by the Neat1_1/2 probe were dramatically reduced in the salivary gland. None of these tissues exhibited histological abnormalities, as revealed by hematoxylin-eosin staining (HE). Dotted boxes indicate the area shown in the insets at higher magnification. Scale bars, 100 µm. (D) Quantification of Malat1 expression in the brains of wild-type and 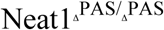 (ΔPAS/ΔPAS) mice (n = 2) using qPCR. The mean values from technical duplicates were used for the statistical analyses. The dots and bars represent the mean value and the standard deviation for the biological duplicates, respectively. p values were calculated by Student’s t-tests.

**Figure 4.**
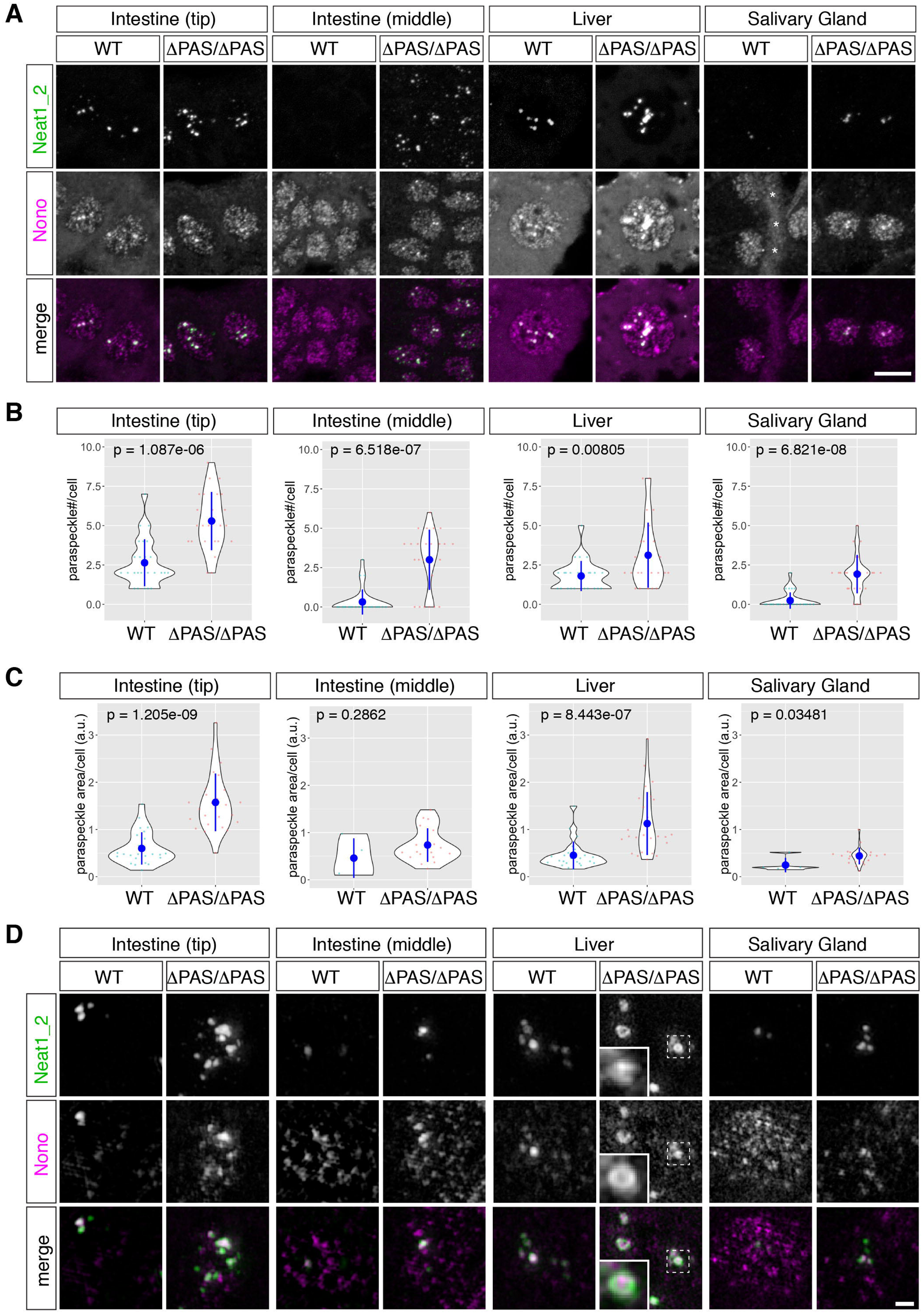
Simultaneous detection of Neat1_2 and the paraspeckle marker Nono in 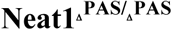 mice. (A) Expression of Neat1_2 in the tip cells of the intestinal epithelium (top), the middle regions of the intestinal epithelium (middle), the liver, and the salivary gland, as revealed by probes that specifically detected Neat1_2 and by the simultaneous detection of the paraspeckle marker Nono. Note the marked increase in paraspeckle formation, which was identified by accumulation of Nono, in the tip cells of the intestine epithelium and the liver cells in 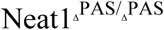 (ΔPAS/ΔPAS) mice. Asterisks in the wild-type salivary gland indicate autofluorescent signals derived from the basement membrane. (B, C) Quantification of the number (B) and the areas (C) of paraspeckles per cell in the tip cells in the intestinal epithelium, the middle of the intestinal epithelium, the liver, and granular duct cells in the salivary gland. Note that the majority of the granular duct cells did not form paraspeckles in the wild-type (WT) mice. Twenty-five cells were randomly selected for the measurement from each tissue derived from two induvial animals for each genotype. p values were calculated by Wilcoxon’s rank sum tests. Blue dots and bars represent the mean value and the standard deviation, respectively. (D) Simultaneous detection of Neat1_2 and Nono at high magnification using structural illumination microscopy. Dotted boxes indicate the area shown in the insets at higher magnification. Scale bars, 10 µm in A and B and 1 µm in D.

Because Neat1 expression was downregulated in Malat1 KO mice (Nakagawa et al. 2012), we examined the expression of Malat1 in 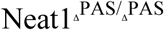 mice to investigate whether these two noncoding RNAs reciprocally regulated one another (**Figure 3D**). The expression of Malat1 was not significantly altered in the liver, intestine, and salivary gland of 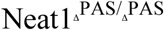 mice (n = 3), consistent with previous observations that Malat1 expression was not affected in Neat1 KO mice (Nakagawa et al. 2011).

To further investigate the formation of paraspeckles at a high resolution, we observed paraspeckles using structured illumination microscopy (**Figure 4D**). The previously described core-shell structure of the paraspeckles (West et al. 2016) was not evident in the tissue sections, except for the liver sections of 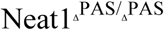 mice, probably because of the freeze-thaw process that was performed during the preparation of the samples (**Figure 4D**). However, paraspeckle formations were clearly recognizable by the accumulation of Neat1_2 and Nono, which formed spherical structures in the cells that strongly expressed Neat1_2 (**Figure 4D**). In the salivary glands of 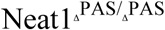 mice, Neat1_2 overlapped with Nono but did not form distinct paraspeckles with round shapes (**Figure 4D**), suggesting that the assembly of Neat1 ribonucleoproteins was incomplete in this cell type. In the livers of 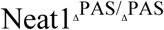 mice, we observed core-shell structures (insets in **Figure 4D)** that were typically observed with cultured cells (West et al. 2016). Given that the liver cells from WT mice expressed only Neat1_2 (**Figure 2B**), upregulated Neat1_2 might be more stable compared to that in other cells, resulting in the formation of “rigid” paraspeckles that are resistant to the freeze-thaw process during the preparation of tissue sections.

In summary, Neat1_2 was differentially upregulated in the 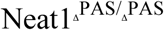 mice, which did not correlate with the expression levels of Neat1_1, resulting in a variable increase in paraspeckle formation in each cell type.

### Cellular differentiation of the intestinal epithelium and salivary gland was not altered in the 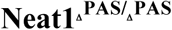 and Neat1 KO mice

Finally, we examined whether the molecular markers of cellular differentiation were altered in tissues that exhibited substantial changes in Neat1 expression in 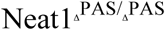 mice. For these studies, we initially focused on the intestinal epithelium, where the ectopic formation of paraspeckles was clearly observed in the middle and bottom regions of the intestinal epithelium. The intestinal epithelium is composed of the following 5 different cell types, which can be identified by the expression of marker genes: multipotent stem cells that produce all of the cells in the intestinal epithelium (*Lgr5*+ cells), Paneth cells that produce antimicrobial peptides (*Lyz1*+ cells), goblet cells that secrete mucous (*Muc2*+ cells), enteroendocrine cells that secrete intestinal hormones (*Chga*+ cells), and enterocytes that are involved in nutrient uptake (Montgomery and Breault 2008) (**Figure 5A**). Enterocytes change their molecular properties as they are displaced towards the top of microvilli; bottom cells express high levels of *Cyb5r3* and *Pigr*, tip cells express high levels of *Apob*, and the topmost cells express *Nt5e* (Moor et al. 2018) (**Figure 5A**). All of these molecular markers in 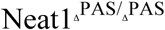 mice exhibited similar expression patterns to those in WT mice (**Figure 5B**), suggesting that the organogenesis of the intestinal epithelium was not affected by the hyperformation of paraspeckles. We also examined the salivary gland, which strongly expressed Neat1_1, and found that Neat1_1 expression was completely lost in the salivary gland of mutant mice. The murine salivary gland is mainly composed of two cell types: acinar cells (Nkcc1+ cells) that secrete saliva (Evans et al. 2000) and granular duct cells that produce many growth factors and hormones, including NGF (Schwab et al. 1976) (**Figure 5C**). Again, we did not observe significant changes in the expression of these molecular markers in 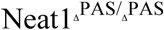 mice (**Figure 5D**). Together, these results and the results of histological analyses using HE staining indicated that the deletion of the Neat1 PAS did not lead to observable phenotypes despite causing the hyperformation of paraspeckles and the loss of Neat1_1 expression. We also examined the expression of intestinal zonation markers and salivary gland markers in Neat1 full KO mice, which essentially lack the expression of both Neat1_1 and Neat1_2 (Nakagawa et al. 2014). We did not observe any recognizable changes in marker expression, suggesting that the cellular differentiation of the intestine and salivary gland is not regulated by the formation of paraspeckles during normal development (**Supplemental Figure S4**).

**Figure 5.**
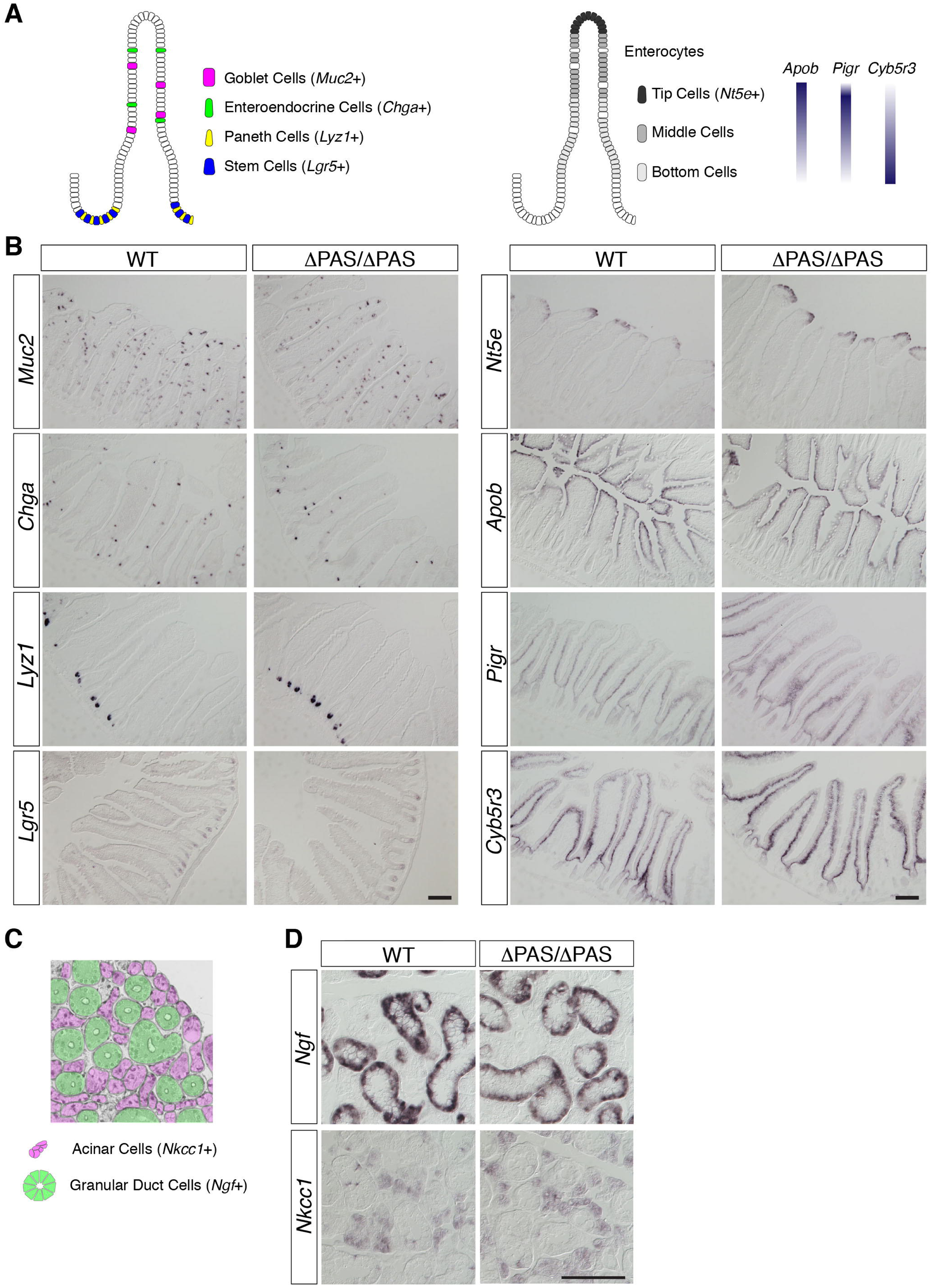
The expression patterns of marker genes were not significantly altered in the intestine and salivary gland of 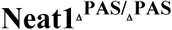 mice. (A) Schematics of the expression patterns of marker genes expressed in the intestinal epithelium. Note that the zonation of the enterocytes can be distinguished by different combinations of the marker genes. (B) In situ hybridization analyses of marker genes in the intestine. (E) Schematics of the expression pattern of the acinar cell marker *Nkcc1* and the granular duct cell marker *Ngf*. (F) In situ hybridization analyses of marker genes in the salivary gland. Scale bars, 100 µm.

## Discussion

Here, we demonstrated that forced isoform switching by deletion of the Neat1 PAS leads to the loss of Neat1_1 and the concomitant upregulation of Neat1_2 in various mouse tissues, as was previously reported for U2OS cells (Li et al. 2017), HAP1 cells (Yamazaki et al. 2018), and ES cells (Modic et al. 2019). The ectopic formation of paraspeckles, as confirmed by the colocalization of Neat1_2 and the paraspeckle marker protein Nono, was observed in cells that normally do not express Neat1_2, including the middle cells in the intestinal epithelium. Enhanced formation of paraspeckles was also observed in cells that endogenously express Neat1_2, including the tip cells in the intestinal epithelium and liver cells. Despite the hyperformation of paraspeckles in various cell types, 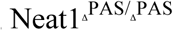 mice were born at the expected Mendelian ratios and did not exhibit any obvious external or histological abnormalities. We also failed to detect any defects in cellular differentiation in 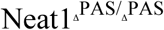 mice using the selected molecular markers. These observations are inconsistent with a recent result showing that hyperformation of paraspeckles upon deletion of the Neat1 PAS in mouse ES cells leads to enhanced expression of differentiation marker genes in tetraploid complemented embryos (Modic et al. 2019). However, the morphological appearance of these embryos was comparable to that of the control embryos reconstituted with WT ES cells, suggesting that enhanced formation of paraspeckles does not interfere with normal development. Thus, the expression of specific genes may be affected in 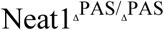 mice, but we failed to detect these changes because we used a limited number of marker genes. This hypothesis is consistent with a recent report showing that deletion of PAS in cultured cell lines leads to only minor changes in gene expression patterns and does not result in defects in cellular proliferation or cell division (Adriaens et al. https://doi.org/10.1101/574582). Alternatively, tetraploid complementation provides a non-natural environment where paraspeckles exert their molecular functions. This idea is supported by the lack of observable embryonic abnormalities in Neat1 KO mice, whereas embryos reconstituted with ES cells lacking Neat1_2 expression exhibit abnormal axis formation and morphological defects. This finding is consistent with the hypothesis that paraspeckles play essential roles when cells are placed under particular, and likely stressful, conditions and are dispensable in normal laboratory conditions (Nakagawa et al. 2011; Nakagawa et al. 2014). Indeed, the expression of Neat1_2 and paraspeckle formation are enhanced by various stimuli, including viral infection (Saha et al. 2006), prolonged cell culture (Nakagawa et al. 2011), proteasome inhibition (Hirose et al. 2014), poly-I:C transfection (Imamura et al. 2014), and mitochondrial stress (Wang et al. 2018). Importantly, Neat1 is controlled by the p53 pathway, but cancer progression is enhanced or inhibited in a context-dependent manner: the lack of Neat1 enhanced the development of premalignant pancreatic intraepithelial neoplasias in KrasG12D-expressing mice (Mello et al. 2017), whereas the progression of skin neoplasias induced by carcinogens was suppressed in Neat1 KO mice (Adriaens et al. 2016). Therefore, 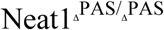 mice may exhibit certain phenotypes when the animals are placed under appropriate experimental conditions.

While previous studies demonstrated that the alternative use of the PAS results in isoform switching of Neat1_1 and Neat1_2 (Naganuma et al. 2012; Li et al. 2017; Yamazaki et al. 2018; Modic et al. 2019), we observed that the upregulation of Neat1_2 in 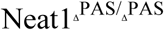 mice was highly variable among the tissues and did not correlate with the expression level of Neat1_1. For example, granular duct cells highly expressed Neat1_1, which should result in the abundant production of Neat1_2 in 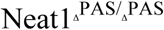 mice. However, Neat1_2 was upregulated only moderately in these cells. As granular duct cells express low levels of Fus (S.N., unpublished observations), which is essential for the assembly of Neat1 ribonucleoprotein complexes into paraspeckles (Naganuma et al. 2012; Shelkovnikova et al. 2014; West et al. 2016), the induced Neat1_2 might show low stability in this cell type. However, Neat1_2 was upregulated in the liver of 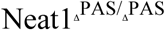 mice despite a lack of detectable expression of Neat1_1 on Northern blots, which cannot be explained by the simple isoform switching mechanism. We speculate that Neat1_1 is produced in liver cells but that it is unstable and is rapidly degraded, and the deletion of PAS may convert these cryptic Neat1_1 transcripts into stable Neat1_2, resulting in the enhanced formation of paraspeckles. Considering that Hnrnpk and Cpsf6 are ubiquitously expressed in mouse tissues, the ratio of Neat1_1 and Neat1_2 might be regulated by the differential stabilization of each isoform in certain cell types. Alternatively, post-translational modification of Hnrnpk and Cpsf6 and/or the isoforms of these proteins might differentially affect the selection of the Neat1 PAS. Thus, researchers should investigate the post-transcriptional regulation of Neat1 transcripts and the precise molecular features of the associated proteins to fully elucidate the physiological roles of paraspeckles and the molecular mechanisms that lead to the differential formation of paraspeckles in living organisms.

In many mouse tissues, Neat1_1 is much more abundant than Neat1_2, and its sequence, including the poly-A signal, is evolutionarily conserved between species. In addition, Neat1_1 localizes to non-paraspeckle sites and forms “microspeckles” in the cells that do not form paraspeckles (Li et al. 2017), suggesting that Neat1_1 may have paraspeckle-independent functions. Because 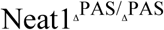 mice lack Neat1_1 expression, they can serve as a model to address the specific functions of Neat1_1. 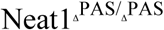 female mice did not exhibit decreased fertility (IM and SN, unpublished observation), suggesting that the impaired formation of corpus luteal cells in the Neat1 KO mice (Nakagawa et al. 2014) was caused by the lack of Neat1_2 but not Neat1_1 expression. The lack of overt phenotypes in 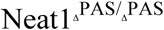 mice also suggests that Neat1_1 does not have an essential role, at least during normal development under laboratory conditions. The evolutionary conservation of Neat1_1 may be explained by the ability of these sequences to stabilize Neat1_2 (Yamazaki et al. 2018), which shares the same sequence. The Neat1 PAS may be conserved because it enables acute induction of Neat1_2 without changes in promotor activities, and Neat1_1 is a by-product of the alternative use of the Neat1 PAS. Identification of a condition where acute isoform switching has an essential role is needed to test this possibility.

## Materials and methods

### Animals

All experiments were approved by the safety division of Hokkaido University (#2015-079). For anesthetization, medetomidine-midazolam-butorphanol (Kawai et al. 2011) was intraperitoneally injected at a volume of 10 µl/g of body weight. The genotyping of Neat1 ΔPAS mice was performed with Quick Taq HS DyeMix (Toyobo #DTM-101, Japan) and the following primers. Neat1_PAS_KO_FW: 5’-GGGAAGCTGTTAAACTGTCAGTC-3’ Neat1_PAS_KO_RV: 5’-TCGGTATACAGGATGCTTTG-3’ The following PCR conditions were used: one cycle at 94°C for 2 min; 35 cycles of 94°C for 30 sec, 57°C for 30 sec, and 68°C for 1 min; followed by 68°C for 5 min. For determination of the Mendelian ratio, heterozygous animals extensively backcrossed to C57BL6/N animals (>8 times) were mated, and the genotype of the offspring was determined at weaning (3 weeks). For statistical analyses, a chi-square test was performed.

### T7 endonuclease assay

For generation of plasmid vectors expressing gRNAs designed to target various regions around the Neat1 PAS, the vectors were amplified with the following forward (F) and reverse (R) primers, and a modified U6-expression vector was used as a template (Inui et al. 2014).

> gRNA#1F: 5’-TCCCCTTTACAGCAAAATAAGTTTTAGAGCTAGAAATAGC-3’
>
> gRNA#1R: 5’-TTATTTTGCTGTAAAGGGGACGGTGTTTCGTCCTTTCCAC-3’
>
> gRNA#2F: 5’-TGCTGTAAAGGGGAGGAAAAGTTTTAGAGCTAGAAATAGC-3’
>
> gRNA#2R: 5’-TTTTCCTCCCCTTTACAGCACGGTGTTTCGTCCTTTCCAC-3’
>
> gRNA#3F: 5’-AGGGGAGGAAAATGGTTAGTGTTTTAGAGCTAGAAATAGC-3’
>
> gRNA#3R: 5’-ACTAACCATTTTCCTCCCCTCGGTGTTTCGTCCTTTCCAC-3’
>
> gRNA#4F: 5’-GAAGCTTCTTAGAATTGTCAGTTTTAGAGCTAGAAATAGC-3’
>
> gRNA#4R: 5’-TGACAATTCTAAGAAGCTTCCGGTGTTTCGTCCTTTCCAC-3’
>
> gRNA#5F: 5’-AAACCTTTATTTTGCTGTAAGTTTTAGAGCTAGAAATAGC-3’
>
> gRNA#5R: 5’-TTACAGCAAAATAAAGGTTTCGGTGTTTCGTCCTTTCCAC-3’

After sequence validation, 3T3 cells were transfected with the gRNA-expressing plasmids and the Cas9-expressing plasmid hCas9 (Addgene #41815) using Lipofectamine LTX reagent (Thermo Fisher #15338100) following the manufacturer’s instructions. Genomic DNA was isolated using a standard procedure 65 hours after the transfection, and the DNA fragments around the target sequences were amplified with the following primers using PrimeSTAR Max (TaKaRa, Japan).

> Neat1_T7_FW: 5’-GGGAAGCTGTTAAACTGTCAGTC-3’
>
> Neat1_T7_RV: 5’-GACAGAATTGCATGTACAGACTG-3’

The following PCR conditions were used: 1 cycle at 98°C for 1 min; followed by 30 cycles of 10 sec at 94°C, 10 sec at 55°C, and 20 sec at 72°C; and one cycle at 72°C for 2 min.

Amplified fragments were purified using a Wizard SV Gel and PCR Clean-Up System (Promega #A9281), diluted at a concentration of 1 µg/20 µl, and subjected to denaturation-denaturation reactions with the following conditions: 1 cycle at 95°C for 1 min and 60 cycles at 85°C for 10 sec with a 1°C decrease for each cycle; the samples were then held at 25°C. The reannealed fragments were treated with 5 units of T7 endonuclease 1 (NEB, #M0302S) at 37°C for 25 min and immediately separated with 7% polyacrylamide gel electrophoresis.

### Generation of genome-edited mice using RNA injection

For preparation of template DNA for in vitro-transcribed gRNA, PCR was performed using the following primers, and the gRNA expression vector used for the T7 assay was the template.

Neat1_gRNA1_T7_temp FW:

> 5’-TAATACGACTCACTATAGGGGTCCCCTTTACAGCAAAATAA-3’
>
> gRNA_Vec_RV:
>
> 5’-AAAAAAGCACCGACTCGGTG-3’

The following PCR conditions were used: 1 cycle at 94°C for 2 min; followed by 30 cycles of 98°C for 10 sec and 68°C for 10 sec; and 1 cycle at 68°C for 5 min. Amplified fragments were purified with a QIAquick PCR Purification Kit (QIAGEN, #28104) and subjected to in vitro transcription using an mMESSAGE mMACHINE T7 Transcription Kit (Thermo Fisher #AM1344) following the manufacturer’s instructions. Transcribed RNAs were purified with a MEGAclea Transcription Clean-Up Kit (Thermo Fisher #AM1908). For preparation of Cas9 mRNA, hCas9 was digested with NotI and purified with QIAquick PCR Purification Kit. In vitro transcription was performed using 1 µg of the template plasmid and an mMESSAGE mMACHINE T7 Transcription Kit, and the template plasmid was digested with Turbo DNase (Thermo Fisher #AM2239). Transcribed mRNAs were purified LiCl precipitation and were used for injection. Single-stranded oligo-DNA was synthesized to introduce the EcoRV restriction site. Neat1_EcoRV_ssDNA:

> 5’-TCGCACACGCTTCTCTGTACTAAAGTGTTCAGTGTACAAACCCACTAACC
>
> ATTTTCCTCCCCTTTACAGCAAGATATCGGTTTGAGATTGAAGCTTCTTAGA
>
> ATTGTCATGGCTGTTACATTCTCGACCTCTGTCACTTTGCCTTCATAT-3’

The RNA and DNA samples were mixed with 25 ng/µl Cas9 mRNA, 12.5 ng/µl gRNA, and 100 ng/µl ssDNA-oligo, and the injection into fertilized eggs was performed by RIKEN RRC. For identification of genome-edited mice, genomic DNA was obtained from the tail of the mice using standard procedures, and PCR fragments were obtained using the primers for the T7 nuclease assay and subjected to agarose gel electrophoresis after digestion with EcoRV. The sequences of the amplified DNA fragments that exhibited differential migration were determined using direct sequencing.

### qPCR analyses of Neat1_1/2 and Neat1_2 expression

RNA was obtained from mouse tissues using TRIzol reagent (Thermo Fisher #15596026) following the manufacturer’s instructions. For increased recovery of Neat1_2, homogenized lysates were incubated at 55°C for 5 min, as previously described (Chujo et al. 2017). After DNase I treatment (Recombinant DNase I, TaKaRa #2270A), cDNAs were synthesized with ReverTra Ace qPCR RT Master Mix (TOYOBO, #FSQ201), and qPCR was performed with a TaKaRa Thermal Cycler Dice TP870 using TB Green Premix Ex TaqII (TaKaRa #RR820A) and the following primers.

> qPCR_Gapdh_FW: 5’-CCTCGTCCCGTAGACAAAATG-3’
>
> qPCR_Gapdh_RV: 5’-TCTCCACTTTGCCACTGCAA-3’
>
> qPCR_Neat1_1/2_FW: 5’-TTGGGACAGTGGACGTGTG-3’
>
> qPCR_Neat1_1/2_RV: 5’-TCAAGTGCCAGCAGACAGCA-3’
>
> qPCR_Neat1_2_FW: 5’-GCTCTGGGACCTTCGTGACTCT-3’
>
> qPCR_Neat1_2_RV: 5’-CTGCCTTGGCTTGGAAATGTAA-3’

### Probe preparations for in situ hybridization

Neat1_1 and Neat1_2 were detected using probes that were described previously (West et al. 2016). For other markers, DNA fragments were prepared by RT-PCR as follows. For preparation of cDNAs, 1 µg of total RNA was reverse transcribed using ReverTra Ace (TOYOBO #TRT-101) and oligo-dT (TOYOBO, #FSK-201) or random primers (TOYOBO #FSK-301) at 30°C for 10 min, followed by 42°C for 45 min, 50°C for 15 min, and 99°C for 5 min. Mixed cDNAs were used as a template for the subsequent PCR using PrimeSTAR Max DNA Polymerase (TaKaRa #R045A). The following PCR conditions were used: 1 cycle at 98°C for 1 min; 35 cycles at 98°C for 10 sec, 55°C for 5 sec, and 72°C for 10 sec; followed by 72°C for 1 min. Amplified fragments were purified using a Wizard SV Gel and PCR Clean-Up System and subcloned into pCRII using a TOPO TA Cloning™ Kit Dual Promoter (Thermo Fisher, #450640). The following primers were used.

> Chga_FW: 5’-ACACAGCCACCAATACCCAA-3’
>
> Chga_RV: 5’-GTTATTGCAGTTGTGCCCCA-3’
>
> Muc2_FW: 5’-GACCTGACAATGTGCCCAGA-3’
>
> Muc2_RV: 5’-AGAAAACGGGGGCTAGAACG-3’
>
> Lyz1_FW: 5’-TCGTTGTGAGTTGGCCAGAA-3’
>
> Lyz1_RV: 5’-AGGCACAGCTCACTAGTCCT-3’
>
> Lgr5_FW: 5’-ATGCGTTTTCTACGTTGCCG-3’
>
> Lgr5_RV: 5’-GGTTGACTCACAGGACCGTT-3’
>
> Nt5e_FW: 5’-CGATCGTCTACCTGGATGGC-3’
>
> Nt5e_RV: 5’-GTTTCCCCTTACCCACTCCC-3’
>
> Apob_FW: 5’-TCCAGAGAGTAGCCCGTGAT-3’
>
> Apob _RV: 5’-AGTTCAGGCTGCTTTGAAGGT-3’
>
> Pigr_FW: 5’-ATCTTCTGTGAGCTGCGACC-3’
>
> Pigr_RV: 5’-CGTAACTAGGCCAGGCTTCC-3’
>
> Cyb5r3_FW: 5’-GGTTGTTGGGGGAGCGG-3’
>
> Cyb5r3_RV: 5’-GAGTTGGAAGGTGGTCCCTG-3’
>
> Ngf_FW: 5’-GAGTTTTGGCCTGTGGTC-3’
>
> Ngf_RV: 5’-TCGGTATACAGGATGCTTTG-3’
>
> Nkcc1_FW: 5’-AGGTCTCTCTGTCGTCGTAA-3’
>
> Nkcc1_RV: 5’-ATGGCCACAGATCATTAAAC-3’

For preparation of a template for in vitro transcription, DNA fragments were amplified by PCR using the M13 FW and RV primers and cDNAs that were subcloned into pCRII as a template. Amplified DNAs were purified with a Wizard SV Gel and PCR Clean-Up System and were used for in vitro transcription with Fluorescein RNA labeling Mix (Merck, #11685619910) and T7 (Merck, #10881767001) or SP6 polymerase (Merck, #11487671001).

### In situ hybridization and immunohistochemistry

In situ hybridization was performed as described previously (Nakagawa et al. 2011; Mito et al. 2016). Briefly, anesthetized adult mice were perfused with 4% paraformaldehyde (PFA) in PBS. Dissected tissues were further fixed for 16 hours at 4°C, rinsed in PBS, and cryoprotected in 30% sucrose in PBS. After the samples were embedded in Tissue-Tek OCT compound (Sakura, Japan), frozen sections were cut at a thickness of 8 µm and were collected on poly-L-lysine (PLL)-coated slide glasses (Matsunami glass #S7441). After rehydration in PBS, the sections were subjected to in situ hybridization with Proteinase K treatment (Mito et al. 2016). MEF cells were cultured on PLL-coated coverslips and fixed in 4% PFA in PBS, rinsed in PBS, permeabilized in 0.5% Triton X-100 for 10 min at room temperature, and processed for in situ hybridization without Proteinase K treatment (Mito et al. 2016). For immunohistochemistry, mice were perfused with 10 ml of PBS to reduce background signals from the endogenous IgGs, and dissected tissues were freshly frozen in OCT compound. Sections were fixed in 4% PFA overnight at 4°C, and antigen retrieval was performed in HistoVT One (Nacalai, Japan, #06380) for 20 min in a Coplin jar placed in boiling water. The antigen retrieval treatment strongly reduces background signals derived from endogenous mouse IgG when using mouse monoclonal antibodies. The following antibodies were used: mouse monoclonal anti-Sfpq (Abcam, #ab11825), mouse monoclonal anti-Nono (Souquere et al. 2010), mouse monoclonal anti-Pspc1 (clone 1L4, Sigma), mouse monoclonal anti-Fus (clone 4H11, Santa Cruz), rabbit polyclonal anti-Brg1 (#A300-813A, Bethyl Laboratories), rabbit polyclonal anti-Tardbp (#10782-2-AP, Proteintech), Cy3-conjugated goat anti-mouse IgG (Merck, #AP124C), Alexa 488-conjugated anti-rabbit IgG (Invitrogen, #A11029), and rabbit polyclonal anti-FITC (Abcam, #ab73831). For preparation of paraffin-embedded sections, perfused tissues were postfixed in Bouin’s solution (Merck, #HT10132) for 16 hours at 4°C and were processed using standard procedures. The bright-field images were obtained with an epifluorescence microscope (BX51; Olympus) equipped with a CCD camera (DP70). Fluorescent and structural illumination microscopy images were obtained with a Zeiss PS1 microscope with 63X water immersion objectives. Fluorescent images of tissue sections were obtained with a Zeiss LSM800 confocal microscope equipped with 63X water immersion objectives. The quantification of paraspeckle areas was performed by measuring the Neat1_2-positive areas that overlapped with Nono signals using ImageJ software.

### Quantification of paraspeckle formations

For quantification of the number and areas of paraspeckles, signals above the background signals were binarized using “Image> Adjust> Threshold” of ImageJ software. Segmentation was performed using “Process> Binary> Watershed”, and the area and the number of each segment were calculated using “Analyze> Analyze particles”. Statistical analyses were performed using R.

### Northern blotting

Five micrograms of total RNA was separated on a denaturing gel (1% agarose gel, 1X MOPS, 2% formaldehyde), the gels were rinsed twice in distilled water for 15 min, and the RNA samples were hydrolyzed in 0.05N NaOH for 20 min. This alkali treatment step is critical for detecting Neat1_2 transcripts. After the samples were soaked in water and subsequently incubated twice in 20X SSC for 45 min, the RNA samples were transferred to a positively charged nylon membrane (Merck, #11209299001) and were hybridized with probes at a concentration of 1 µg/ml in DIG Easy Hyb (Merck, #11603558001). The hybridized probes were detected with anti-Dig AP (Merck, #11093274910) and CDP-Star (Merck, #11685627001) following the manufacturer’s instructions.

### Western blotting

Five micrograms of total protein was separated on a precast 10–20% gradient gel (Funakoshi, #291M) and blotted on a nitrocellulose membrane using a semi-dry blotter following a standard protocol. Antibodies were diluted in Can Get Signal Immunoreaction Enhancer Solution (Toyobo, NKB-101) and detected with ECL Prime (GE Healthcare, #RPN2232).

## Acknowledgments

This work was supported by JSPS KAKENHI Grant Numbers 17H03604, 16H06279, and 16H06276, MEXT KAKENHI Grant Number 26113005, a Toray Science and Technology Grant to S.N., and a grant of the Joint Research Program of IGM, Hokkaido University to S.N. and T.H. We thank Dr. Yuko Okamatsu for the use of the confocal microscope.

## Author contributions

M.I. and H.T. performed the immunostaining, probe preparation, and in situ hybridization, M.M., T.C., and H.A. designed and created the mutant mice, T.H. and S.N. designed the experiments, and S.N. wrote the paper with contributions from all authors.

## References

Adriaens C, Standaert L, Barra J, Latil M, Verfaillie A, Kalev P, Boeckx B, Wijnhoven PW, Radaelli E, Vermi W et al. 2016. p53 induces formation of NEAT1 lncRNA-containing paraspeckles that modulate replication stress response and chemosensitivity. Nat Med 22: 861–868.

Ahmed ASI, Dong K, Liu J, Wen T, Yu L, Xu F, Kang X, Osman I, Hu G, Bunting KM et al. 2018. Long noncoding RNA NEAT1 (nuclear paraspeckle assembly transcript 1) is critical for phenotypic switching of vascular smooth muscle cells. Proc Natl Acad Sci U S A 115: E8660–E8667.

Chen LL, Carmichael GG. 2009. Altered nuclear retention of mRNAs containing inverted repeats in human embryonic stem cells: functional role of a nuclear noncoding RNA. Mol Cell 35: 467–478.

Chujo T, Yamazaki T, Kawaguchi T, Kurosaka S, Takumi T, Nakagawa S, Hirose T. 2017. Unusual semi-extractability as a hallmark of nuclear body-associated architectural noncoding RNAs. EMBO J 36: 1447–1462.

Clemson CM, Hutchinson JN, Sara SA, Ensminger AW, Fox AH, Chess A, Lawrence JB. 2009. An architectural role for a nuclear noncoding RNA: NEAT1 RNA is essential for the structure of paraspeckles. Mol Cell 33: 717–726.

Dettwiler S, Aringhieri C, Cardinale S, Keller W, Barabino SM. 2004. Distinct sequence motifs within the 68-kDa subunit of cleavage factor Im mediate RNA binding, protein-protein interactions, and subcellular localization. J Biol Chem 279: 35788–35797.

Evans RL, Park K, Turner RJ, Watson GE, Nguyen HV, Dennett MR, Hand AR, Flagella M, Shull GE, Melvin JE. 2000. Severe impairment of salivation in Na+/K+/2Cl-cotransporter (NKCC1)-deficient mice. J Biol Chem 275: 26720–26726.

Fong KW, Li Y, Wang W, Ma W, Li K, Qi RZ, Liu D, Songyang Z, Chen J. 2013. Whole-genome screening identifies proteins localized to distinct nuclear bodies. J Cell Biol 203: 149–164.

Fox AH, Bond CS, Lamond AI. 2005. P54nrb forms a heterodimer with PSP1 that localizes to paraspeckles in an RNA-dependent manner. Mol Biol Cell 16: 5304–5315.

Fox AH, Lam YW, Leung AK, Lyon CE, Andersen J, Mann M, Lamond AI. 2002. Paraspeckles: a novel nuclear domain. Curr Biol 12: 13–25.

Fox AH, Nakagawa S, Hirose T, Bond CS. 2018. Paraspeckles: Where Long Noncoding RNA Meets Phase Separation. Trends Biochem Sci 43: 124–135.

Hirose T, Virnicchi G, Tanigawa A, Naganuma T, Li R, Kimura H, Yokoi T, Nakagawa S, Benard M, Fox AH et al. 2014. NEAT1 long noncoding RNA regulates transcription via protein sequestration within subnuclear bodies. Mol Biol Cell 25: 169–183.

Hu SB, Xiang JF, Li X, Xu Y, Xue W, Huang M, Wong CC, Sagum CA, Bedford MT, Yang L et al. 2015. Protein arginine methyltransferase CARM1 attenuates the paraspeckle-mediated nuclear retention of mRNAs containing IRAlus. Genes Dev 29: 630–645.

Imamura K, Imamachi N, Akizuki G, Kumakura M, Kawaguchi A, Nagata K, Kato A, Kawaguchi Y, Sato H, Yoneda M et al. 2014. Long noncoding RNA NEAT1-dependent SFPQ relocation from promoter region to paraspeckle mediates IL8 expression upon immune stimuli. Mol Cell 53: 393–406.

Inui M, Miyado M, Igarashi M, Tamano M, Kubo A, Yamashita S, Asahara H, Fukami M, Takada S. 2014. Rapid generation of mouse models with defined point mutations by the CRISPR/Cas9 system. Sci Rep 4: 5396.

Kawaguchi T, Tanigawa A, Naganuma T, Ohkawa Y, Souquere S, Pierron G, Hirose T. 2015. SWI/SNF chromatin-remodeling complexes function in noncoding RNA-dependent assembly of nuclear bodies. Proc Natl Acad Sci U S A 112: 4304–4309.

Kawai S, Takagi Y, Kaneko S, Kurosawa T. 2011. Effect of three types of mixed anesthetic agents alternate to ketamine in mice. Exp Anim 60: 481–487.

Li R, Harvey AR, Hodgetts SI, Fox AH. 2017. Functional dissection of NEAT1 using genome editing reveals substantial localization of the NEAT1_1 isoform outside paraspeckles. RNA 23: 872–881.

Long Y, Wang X, Youmans DT, Cech TR. 2017. How do lncRNAs regulate transcription? Sci Adv 3: eaao2110.

Mello SS, Sinow C, Raj N, Mazur PK, Bieging-Rolett K, Broz DK, Imam JFC, Vogel H, Wood LD, Sage J et al. 2017. Neat1 is a p53-inducible lincRNA essential for transformation suppression. Genes Dev 31: 1095–1108.

Mito M, Kawaguchi T, Hirose T, Nakagawa S. 2016. Simultaneous multicolor detection of RNA and proteins using super-resolution microscopy. Methods 98: 158–165.

Modic M, Grosch M, Rot G, Schirge S, Lepko T, Yamazaki T, Lee FCY, Rusha E, Shaposhnikov D, Palo M et al. 2019. Cross-Regulation between TDP-43 and Paraspeckles Promotes Pluripotency-Differentiation Transition. Mol Cell 74: 951–965 e913.

Montgomery RK, Breault DT. 2008. Small intestinal stem cell markers. J Anat 213: 52–58.

Moor AE, Harnik Y, Ben-Moshe S, Massasa EE, Rozenberg M, Eilam R, Bahar Halpern K, Itzkovitz S. 2018. Spatial Reconstruction of Single Enterocytes Uncovers Broad Zonation along the Intestinal Villus Axis. Cell 175: 1156–1167 e1115.

Naganuma T, Nakagawa S, Tanigawa A, Sasaki YF, Goshima N, Hirose T. 2012. Alternative 3’-end processing of long noncoding RNA initiates construction of nuclear paraspeckles. EMBO J 31: 4020–4034.

Nakagawa S, Ip JY, Shioi G, Tripathi V, Zong X, Hirose T, Prasanth KV. 2012. Malat1 is not an essential component of nuclear speckles in mice. RNA 18: 1487–1499.

Nakagawa S, Naganuma T, Shioi G, Hirose T. 2011. Paraspeckles are subpopulation-specific nuclear bodies that are n16239143 19716791 25792598ot essential in mice. J Cell Biol 193: 31–39.

Nakagawa S, Shimada M, Yanaka K, Mito M, Arai T, Takahashi E, Fujita Y, Fujimori T, Standaert L, Marine JC et al. 2014. The lncRNA Neat1 is required for corpus luteum formation and the establishment of pregnancy in a subpopulation of mice. Development 141: 4618–4627.

Nakagawa S, Yamazaki T, Hirose T. 2018. Molecular dissection of nuclear paraspeckles: towards understanding the emerging world of the RNP milieu. Open Biol 8.

Saha S, Murthy S, Rangarajan PN. 2006. Identification and characterization of a virus-inducible non-coding RNA in mouse brain. J Gen Virol 87: 1991–1995.

Sasaki YT, Ideue T, Sano M, Mituyama T, Hirose T. 2009. MENepsilon/beta noncoding RNAs are essential for structural integrity of nuclear paraspeckles. Proc Natl Acad Sci U S A 106: 2525–2530.

Schwab ME, Stockel K, Thoenen H. 1976. Immunocytochemical localization of nerve growth factor (NGF) in the submandibular gland of adult mice by light and electron microscopy. Cell Tissue Res 169: 289–299.

Shelkovnikova TA, Robinson HK, Troakes C, Ninkina N, Buchman VL. 2014. Compromised paraspeckle formation as a pathogenic factor in FUSopathies. Hum Mol Genet 23: 2298–2312.

Souquere S, Beauclair G, Harper F, Fox A, Pierron G. 2010. Highly ordered spatial organization of the structural long noncoding NEAT1 RNAs within paraspeckle nuclear bodies. Mol Biol Cell 21: 4020–4027.

Standaert L, Adriaens C, Radaelli E, Van Keymeulen A, Blanpain C, Hirose T, Nakagawa S, Marine JC. 2014. The long noncoding RNA Neat1 is required for mammary gland development and lactation. RNA 20: 1844–1849.

Sunwoo H, Dinger ME, Wilusz JE, Amaral PP, Mattick JS, Spector DL. 2009. MEN epsilon/beta nuclear-retained non-coding RNAs are up-regulated upon muscle differentiation and are essential components of paraspeckles. Genome Res 19: 347–359.

Wang Y, Hu SB, Wang MR, Yao RW, Wu D, Yang L, Chen LL. 2018. Genome-wide screening of NEAT1 regulators reveals cross-regulation between paraspeckles and mitochondria. Nat Cell Biol 20: 1145–1158.

West JA, Mito M, Kurosaka S, Takumi T, Tanegashima C, Chujo T, Yanaka K, Kingston RE, Hirose T, Bond C et al. 2016. Structural, super-resolution microscopy analysis of paraspeckle nuclear body organization. J Cell Biol 214: 817–830.

Wilusz JE, JnBaptiste CK, Lu LY, Kuhn CD, Joshua-Tor L, Sharp PA. 2012. A triple helix stabilizes the 3’ ends of long noncoding RNAs that lack poly(A) tails. Genes Dev 26: 2392–2407.

Yamazaki T, Souquere S, Chujo T, Kobelke S, Chong YS, Fox AH, Bond CS, Nakagawa S, Pierron G, Hirose T. 2018. Functional Domains of NEAT1 Architectural lncRNA Induce Paraspeckle Assembly through Phase Separation. Mol Cell 70: 1038–1053 e1037.

