## Supplemental materials for "Forced isoform switching of Neat1_1 to Neat1_2 leads to the loss of Neat1_1 and the hyperformation of paraspeckles but does not affect the development and growth of mice"

### Figure legends for supplemental materials

**Supplementary Figure S1. Northern blot analyses of Neat1 expression in various tissues of Neat1$^{\Delta PAS/\Delta PAS}$ mice**

(A) Schematics of the probes used for the detection of Neat1_1 and Neat1_2. The Neat1/2 probe detected both isoforms, whereas the Neat1_2 probe targeted a region specific to the long isoform. (B, C) Expression of Neat1 isoforms, as revealed by probes that detected Neat1/2 (B) and Neat1_2. Asterisks indicate lanes with degraded RNAs, which were not included in the statistical analyses shown in **Figure 2D**. (C). Note the variable expression of Neat1_1 and Neat1_2 in the wild-type mice and the variable upregulation of Neat1_2 in Neat1$^{\Delta PAS/\Delta PAS}$ mice.

**Supplementary Figure S2. Northern blot analyses of Neat1 expression in representative tissues of Neat1 KO mice**

(A) Schematics of the probes used for the detection of Neat1_1 and Neat1_2. The Neat1/2 probe detected both isoforms, whereas the Neat1_2 probe targeted a region specific to the long isoform. (B, C) Expression of Neat1 isoforms, as revealed by probes that detected Neat1/2 (B) and Neat1_2 (C).

**Supplementary Figure S3. RT-qPCR analyses of Neat1_2 expression in various tissues of Neat1$^{\Delta PAS/\Delta PAS}$ mice**

(A) Schematics of the regions amplified by RT-qPCR primers used for the detection of Neat1_1/2 and Neat1_2. The Neat1_1/2 primers detected both isoforms, whereas the Neat1_2 primers targeted a region specific to the long isoform. (B) RT-qPCR analyses of Neat1_1/2 and Neat1_2 expression in the RNA samples shown in **Supplemental Figure S2**. The Neat1 expression was normalized by the expression of Gapdh except for liver and colon samples, which were normalized by the expression of β-actin because of variable expression of Gapdh in these tissues. The dots and bars represent the mean value and the standard deviation for the biological triplicates, respectively.

**Supplementary Figure S4. The expression patterns of marker genes were not significantly altered in the intestine and salivary gland of Neat1 KO mice**

(A) Schematics of the expression patterns of marker genes expressed in the intestinal epithelium. Note that the zonation of the enterocytes can be distinguished by different combinations of the marker genes. (B) In situ hybridization analyses of marker genes in the intestine. (E) Schematics of the expression pattern of the acinar cell marker *Nkcc1* and the granular duct cell marker *Ngf*. (F) In situ hybridization analyses of marker genes in the salivary gland. Scale bars, 100 µm.

Isobe et al. Supplemental figure S1

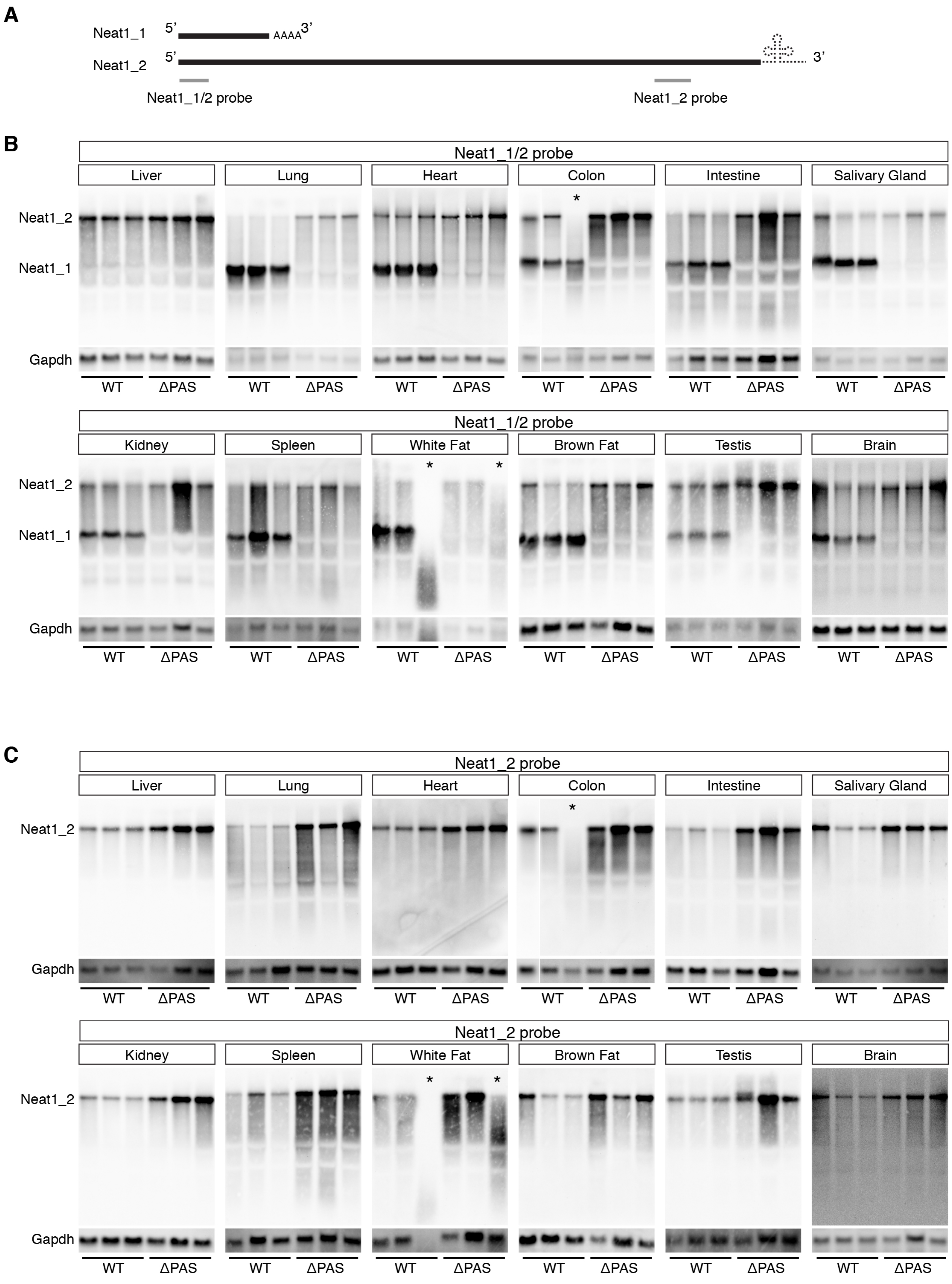

Isobe et al. Supplemental figure S2

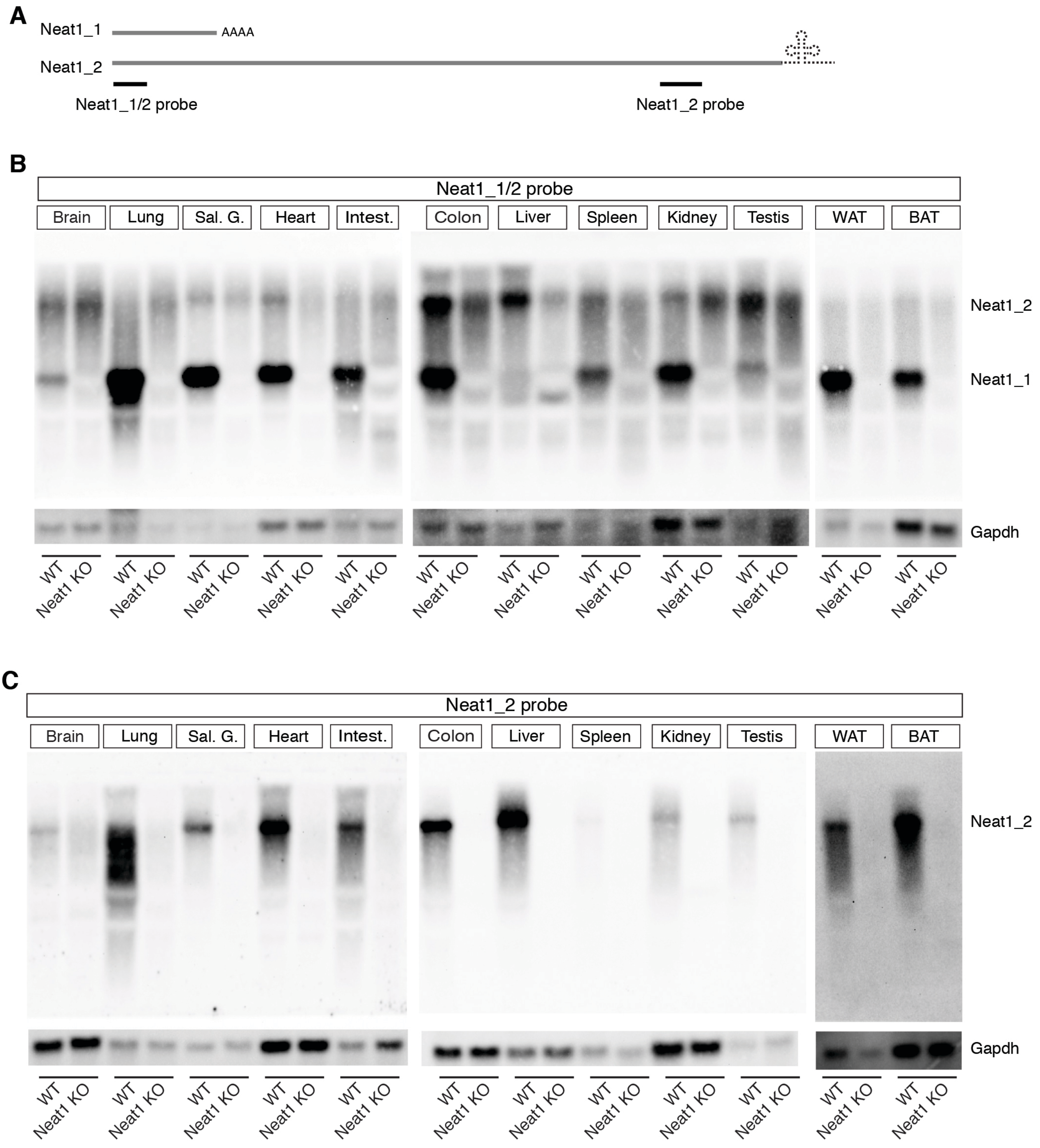

Isobe et al. Supplemental figure S3

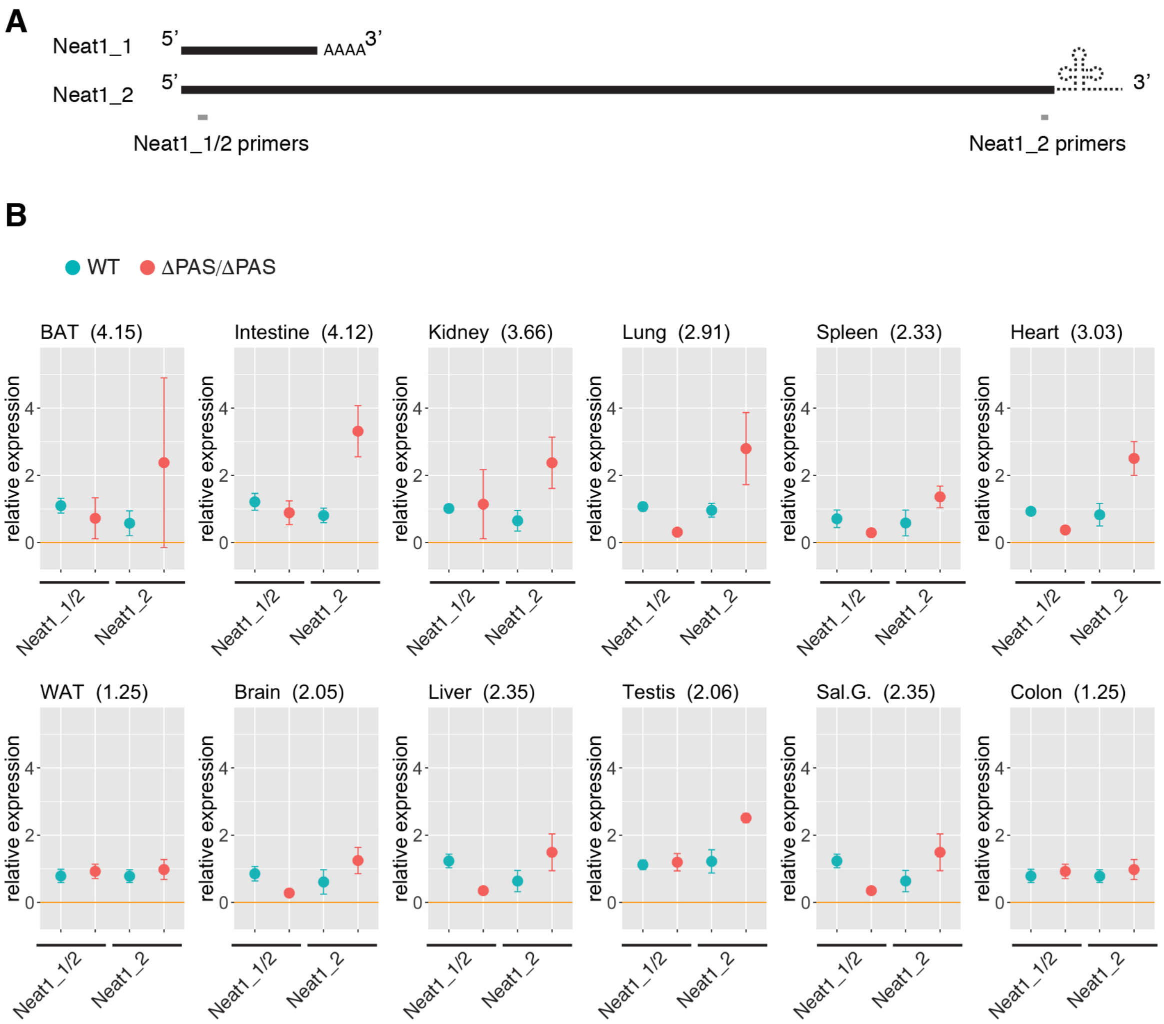

Isobe et al. Supplemental figure S4

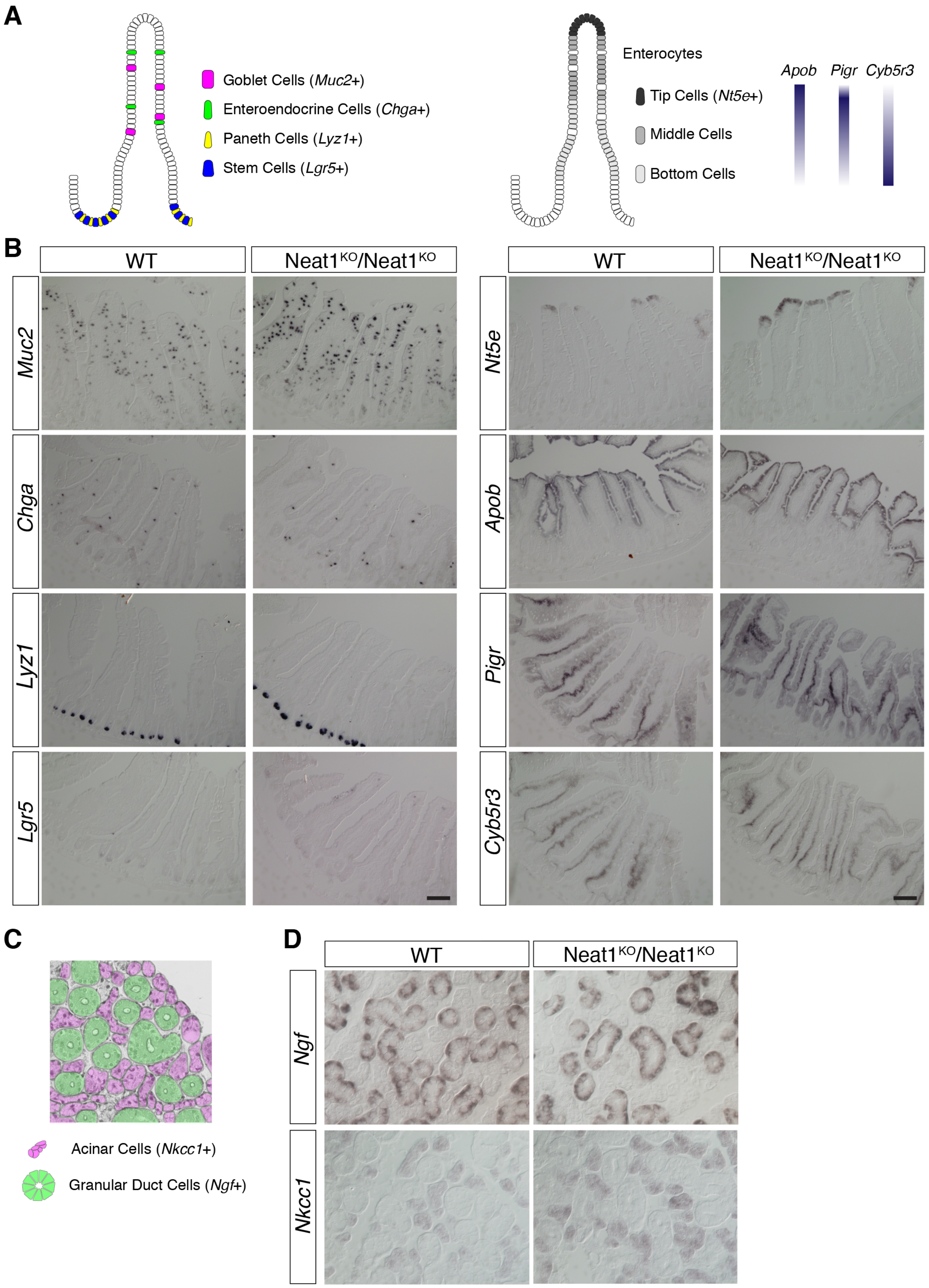
